# SARS-CoV-2 S protein activates the HIV latent reservoir through the mTOR pathway

**DOI:** 10.1101/2025.08.18.670887

**Authors:** Linle Xu, Liangjuan Chen, Hanying Wang, Honglin Shi, Xingzhong Miao, Shiqing Li, Yufen Jiang, Hongbo Shi

## Abstract

During the global COVID-19 pandemic, mRNA vaccines using the S protein as antigen were widely used.Vaccine-induced S proteins can persist in vivo for weeks, triggering low-level immune activation. HIV latent reservoir maintenance is a major challenge for ART therapy, especially when immune pressure is waning. This then raises critical questions for HIV-infected patients: does prolonged exposure to S proteins affect HIV latent reservoir stability? Recent studies have pointed out that S proteins may activate the mTOR signaling pathway, which in turn affects the immune response and metabolic processes of cells. And the mTOR pathway is closely related to the maintenance and activation of HIV latent reservoir. However, how S proteins affect the HIV latent reservoir and the mechanism of activation are unclear. To explore the mechanism of how SARS-CoV-2 S proteins regulate the HIV latent reservoir and to explore whether S proteins regulate the HIV latent reservoir through the mTOR pathway, we constructed an in vitro HIV latent reservoir model for our experiments.To evaluate the potential role of S protein in HIV latent reservoir activation, relevant markers of HIV latent reservoir activation were detected using ELISA, flow cytometry, and RT-qPCR; and the relationship between S protein and mTOR was also detected by WB, CO-IP, and IFC.It was found that S proteins activated the HIV latent reservoir while increasing mTOR expression. It was further observed that mTOR inhibitors significantly inhibited S protein-induced activation of the HIV latent reservoir, and mTOR activators reversed the inhibitory effect of mTOR inhibitors on HIV latent reservoir activation. In summary, we found that S proteins activated the HIV latent reservoir through the mTOR pathway.

S protein interacts with mTOR and activates the mTOR-p-p70S6K-pS6 pathway, which promotes HIV transcription

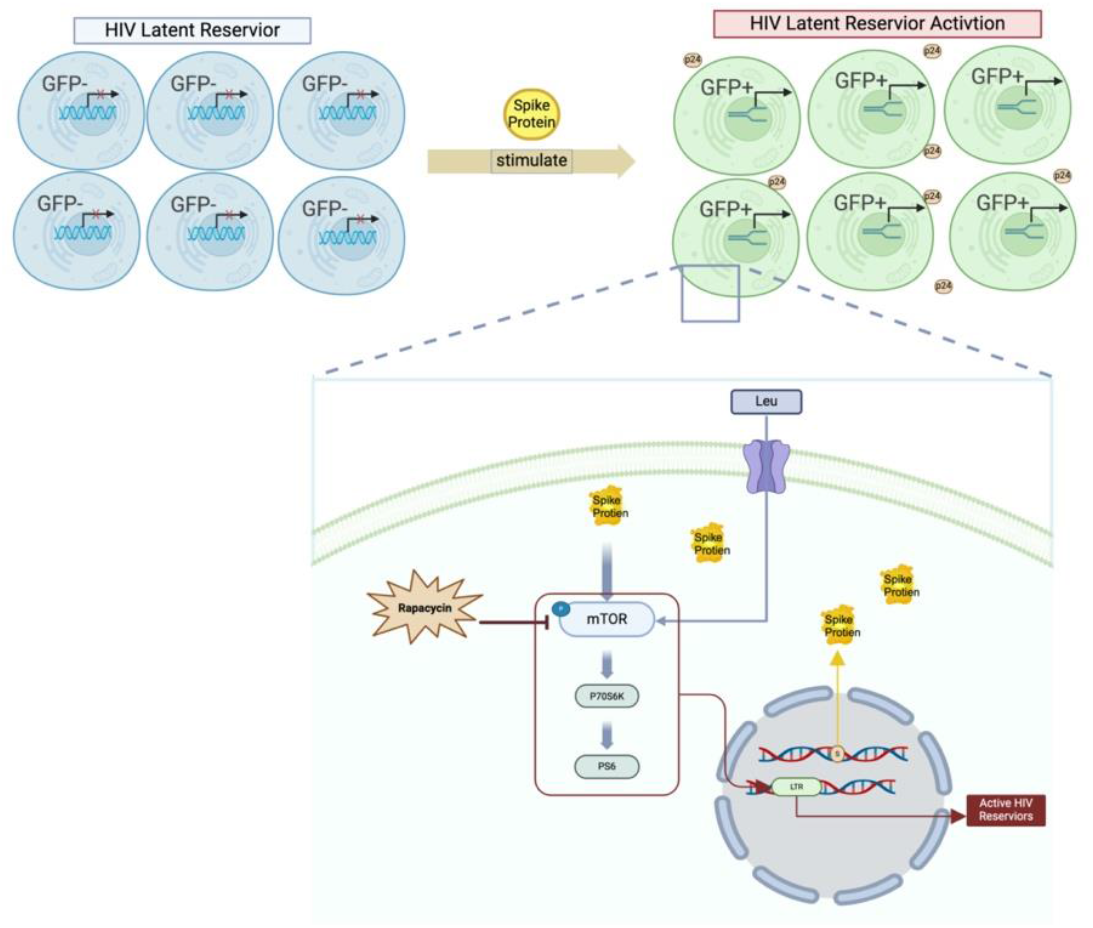

## Introduction

With the global spread of the SARS-CoV-2 epidemic, the development and promotion of COVID-19 vaccines has become a key strategy in global public health. Among them, mRNA vaccines targeting S proteins are widely used in different populations. After vaccination, the body usually undergoes a short-term immune activation, including maturation of dendritic cells (DCs), activation of T and B cells, and release of pro-inflammatory cytokines [1, 2]. In healthy individuals, this immune activation contributes to the induction of a protective immune response, but in HIV-infected individuals, the vaccine-triggered immune response may have additional effects, particularly on the modulation of the HIV latent reservoir [3, 4].Maintenance of the HIV latent reservoir is one of the major challenges facing antiretroviral therapy. Even under long-term antiretroviral therapy, the latently infected HIV genome remains in the body and is stored primarily in resting CD4^+^ T cells and certain myeloid cells.When the immune system is stimulated, these latent viruses may be reactivated, leading to HIV replication and transmission [5, 6]. Therefore, immune activation induced by the COVID-19 vaccine may play an important role in the activation of HIV latent reservoirs [7–9]. A case report describes a 65-year-old HIV patient on ART who developed a transient increase in HIV viral load (VL) 28 days after the first dose of mRNA-1273 vaccine [10].Another study found an increase in HIV viral load after COVID-19 vaccination in older HIV-infected individuals, and possible changes in the diversity of HIV reservoirs and TCR pools in HIV-positive individuals [11].Nevertheless, some studies have found different conclusions. In cohort and population-level analyses, no evidence was found that the COVID-19 mRNA vaccine systematically promotes the release of HIV virus from the reservoir into the plasma [12]. Therefore, after vaccination, the immune system is stimulated to generate an immune response, and whether its effects extend to the HIV latent reservoirs, leading to viral activation, remains a topic of research worth exploring.

The mTOR signaling pathway is an important regulator of cell growth, survival, and immune regulation, and plays a key role in several aspects of cell metabolism, proliferation, and immune response [13, 14].In the context of HIV infection, the activation status of mTOR has an important impact on both the maintenance and activation of the HIV latent reservoir [15, 16].Activation of mTORC1 inhibits autophagy, prolongs the survival of infected cells, and perpetuates the presence of latently infected CD4^+^ T cells, which serve as “shelters” for viral reservoirs [17, 18].T cells in the latent reservoir are usually in a metabolic quiescent state, which is mainly dependent on oxidative phosphorylation, and low mTOR activity may be associated with this state. Inhibition of mTOR may indirectly stabilize the latent reservoir by maintaining cellular quiescence [19];It has also been shown that mTOR activation may provide energy support to latent cells by promoting glycolysis [15].During neocoronary infection, S proteins can interact with host cells via angiotensin-converting enzyme 2 (ACE2) receptors and related co-receptors, which in turn modulate mTOR signaling [20–22].For example, SARS-CoV-2 infection enhances the protein synthesis and metabolic activity of host cells through the PI3K/Akt/mTOR pathway, providing a favorable environment for viral proliferation [23].In addition, studies have shown that mTOR inhibitors are able to reduce the replication capacity of SARS-CoV-2 and inhibit its over-activation of the host immune system to a certain extent [24, 25], suggesting that the role of mTOR in neocollagenic infections deserves to be explored in depth.Given that the S protein of SARS-CoV-2 can activate the mTOR signaling pathway and that mTOR signaling has an important regulatory role in the stability of the HIV latent reservoir, it is of great scientific significance and clinical value to explore whether the S protein can affect the HIV latent reservoir through the mTOR pathway.Based on the clinical observation of transient elevation of viral load in HIV-infected patients after COVID-19 vaccination and the critical dual role of the mTOR signaling pathway in the maintenance of the HIV latent reservoir and SARS-CoV-2 infection, the aim of this study was to investigate whether the SARS-CoV-2 S protein influences the stability and activation status of the HIV latent reservoir by regulating the mTOR signaling pathway. With this study, we hope to reveal potential mechanistic links to changes in the dynamics of the HIV latent reservoir in the context of new coronavirus infection or vaccination.

## Results

### 1. Construction of J-Lat 10.6 cell lines expressing S protein

J-Lat 10.6 cells are a classic HIV latent reservoir model cell line with an integrated green fluorescent protein (GFP) reporting system. When latent HIV viruses are activated, the LTR promoter drives GFP expression, causing the cells to emit green fluorescence, thereby enabling direct assessment of the activation status of the virus. The experiment was conducted using CD3, CD4, and CD8 antibody staining, and a flow cytometer was used to detect cell surface antigen expression. The results showed that J-Lat10.6 cells expressed high levels of CD3 and CD4, while CD8 expression was almost zero, confirming the phenotypic characteristics of this cell line(Figure 1A). Subsequently, J-Lat10.6 cell line stably expressing the SARS-CoV-2 spike protein was established using a lentiviral vector. Western blot was used to validate the expression of S protein.The results indicate that the constructed J-Lat cell line successfully stably expresses the SARS-CoV-2 spike protein (Figure 1B), providing an experimental model for subsequent studies. We use PHA as a T cell activator to break HIV latency. Concentration screening experiments were conducted (concentration range: 1, 2.5, 5, 7.5 μg/mL). Microscopic observation revealed that when the PHA concentration reached 7.5 μg/mL, the cell status was significantly worse (Figure 1C). At concentrations between 1 and 5 μg/mL, the green fluorescent signal gradually intensified with increasing concentration, indicating a gradual increase in activation efficiency (Figure 1D). At the same time, flow cytometry was used to detect GFP expression at various concentrations, and the results were consistent with the fluorescence images (Figure 1E, F). Considering the cell status and activation efficiency, 5 μg/mL was selected as the optimal concentration for subsequent experiments.

**Figure 1:**
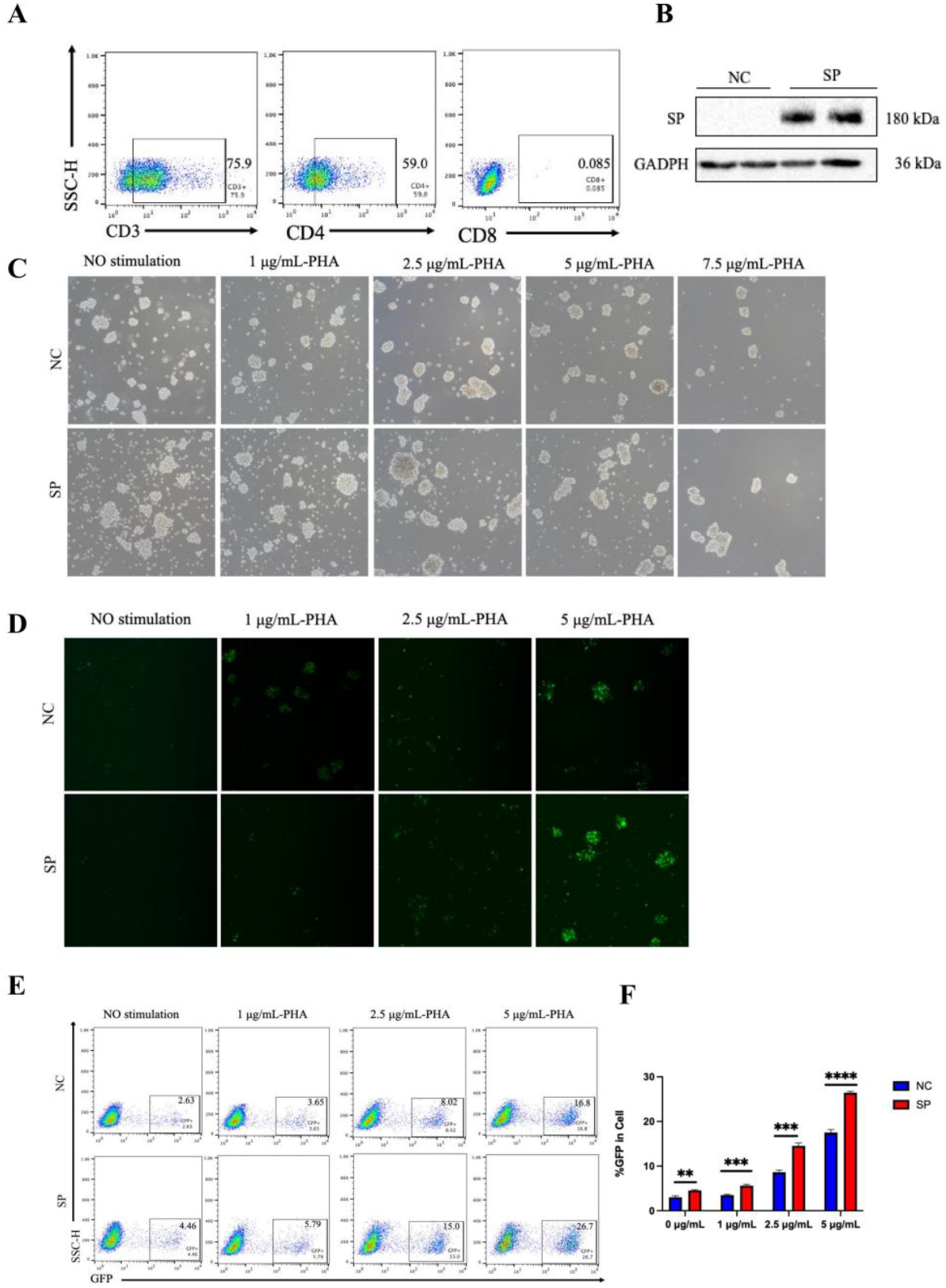
(A) Expression levels and percentage distribution of CD3, CD4, and CD8 molecules on the surface of J-Lat 10.6 cells as detected by flow cytometry. (B) Western blot validation of the successful transfection of the SARS-CoV-2 spike protein into the J-Lat 10.6 cell line. The NC group serves as the empty vector control, and the SP group is the experimental group, with GAPDH as the internal control. (C): Electron micrographs of the concentration screening experiment using different concentrations of PHA (1, 2.5, 5, 7.5 μg/mL). (D): Observation of the effects of different concentrations of PHA on GFP expression in J-Lat 10.6 cells under an inverted fluorescence microscope (magnification: 10×). (E): Percentage of GFP-positive cells in J-Lat 10.6 cells treated with different concentrations of PHA, as determined by flow cytometry. (F) Statistical analysis of the percentage of GFP-positive cells.Data from each experimental group include at least three independent cell experiments, with results expressed as mean ± standard deviation. **p<0.01, ***p<0.001, ****p<0.0001.

### 2. S protein stimulation promotes the activation of the HIV latent reservoir

To comprehensively assess the impact of S protein on HIV latent reservoir activation, this experiment designed multiple control groups and experimental groups. In addition to the empty vector control group (NC) and the S protein transfection group (SP). Two additional groups were also established: one group consisted of the NC group plus PHA (NC+PHA), serving as a known latent reservoir activation positive control to validate the activation efficacy of PHA; the other group was the S protein combined with PHA treatment group (SP+PHA), used to assess whether S protein can synergistically interact with PHA to enhance the activation efficacy of the latent reservoir.

To investigate the effect of S protein on HIV latent reservoir activation, fluorescence inverted microscopy was used to observe electron fluorescence images. Figure 2A shows the changes in GFP signals in cells under different treatment conditions.Flow cytometry was then used to quantitatively analyze GFP expression in cells. The experimental results are shown in Figure 2C. The proportion of GFP-positive cells in the SP group was 16.7%, higher than that in the NC group, indicating that S protein alone can promote the activation of the HIV latent reservoir, thereby increasing the proportion of GFP+cells.RT-qPCR analysis of Gag and LTR gene expression in different treatment groups revealed that, compared with the NC group, Gag and LTR gene expression was significantly upregulated in the SP group, suggesting that the S protein promotes the transcriptional activation of latent viruses. Furthermore, GAG and LTR expression levels in the SP+PHA group were significantly higher than those in the NC+PHA group, indicating that the combination of the S protein and PHA enhances the activation of the latent reservoir (Figure 2DE).To verify the release of particles after viral activation, we measured the levels of P24 antigen in cell culture supernatants using ELISA.The results showed that the P24 levels in the SP group were significantly higher than those in the NC group, further confirming the role of the S protein in latent reservoir activation; simultaneously, the P24 concentrations in the SP+PHA group were significantly higher than those in the NC+PHA group (Figure 2F). This indicates that the combined treatment of the S protein and PHA significantly enhances viral particle release, further supporting the synergistic role of the S protein in latent reservoir activation. To assess changes in the phenotype of HIV latent reservoir cells, flow cytometry was used to detect the expression of CD4 molecules.The results showed that CD4 expression was lower in the SP group than in the NC group, and CD4 expression was lower in the SP+PHA group than in the NC+PHA group, indicating that S protein promotes HIV activation, and the combined treatment of PHA and S protein produced a synergistic effect (Figure 2I). These results suggest that S protein plays an important role in activating the HIV latent reservoir, providing new insights for future studies on the mechanisms of latent reservoir activation.

**Figure 2:**
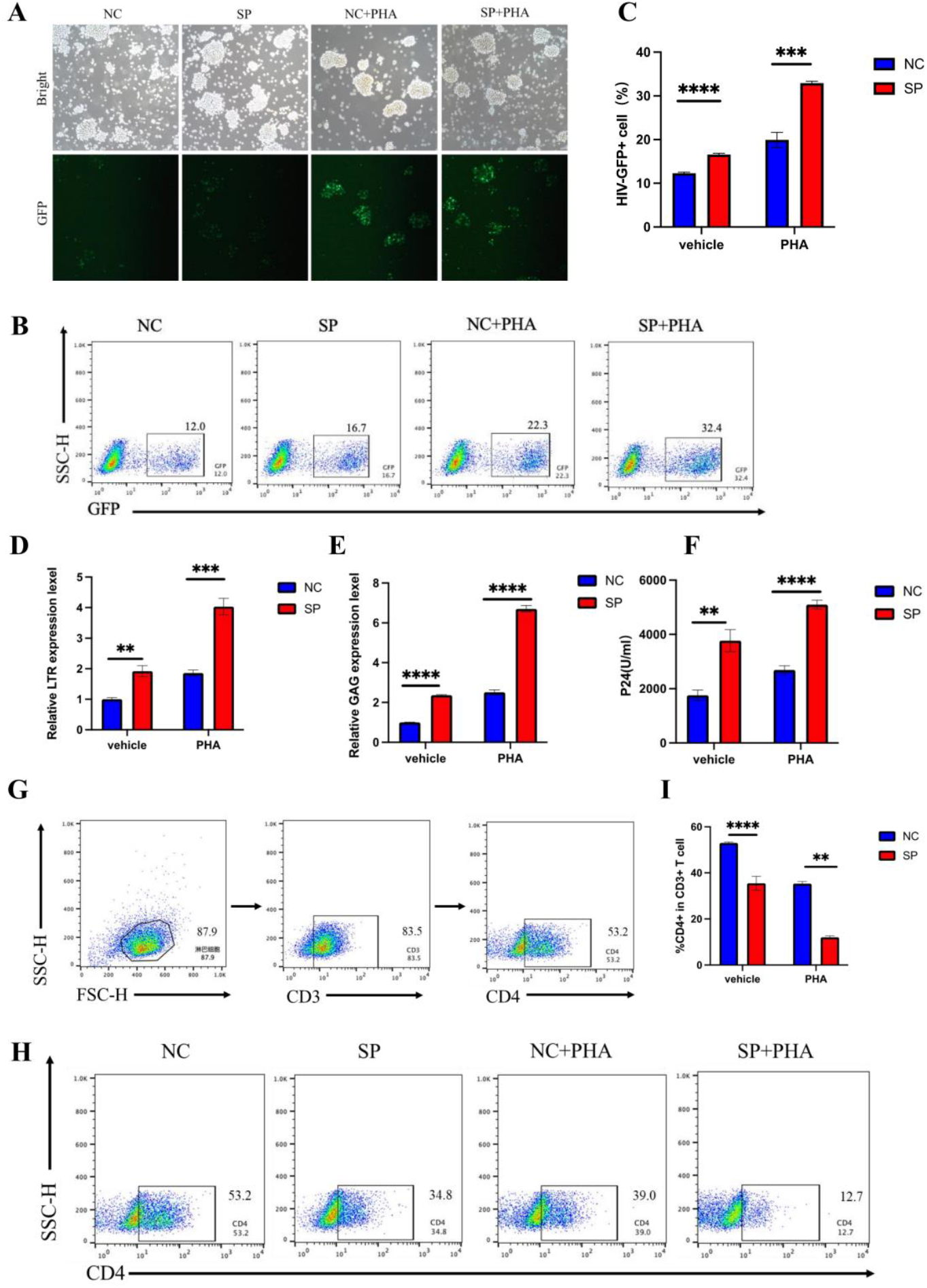
(A) Electron microscope images of cells under different treatments showing GFP expression (magnification: 10×). The NC group is the empty vector control group, the SP group is the S protein-transfected group, the NC+PHA group is the PHA-activated empty vector control group, and the SP+PHA group is the S protein and PHA co-treated group.(B) Representative flow cytometric plots of GFP-positive cells in different groups. (C) Statistical analysis of the percentage of GFP-positive cells. Data from each experimental group include at least three independent cell experiments, with results expressed as mean ± standard deviation. ***p<0.001, ****p<0.0001. (D) Transcriptional levels of HIV LTR in different groups.Data from each experimental group include at least three independent cell experiments, with results expressed as mean ± standard deviation. **p<0.01, ***p<0.001. (E) Transcriptional levels of HIV Gag in different groups. Data from each experimental group include at least three independent cell experiments, with results expressed as mean ± standard deviation. ****p<0.0001. (F) p24 antigen levels in the supernatant of different treatment groups. Data from each experimental group include at least three independent cell experiments, with results expressed as mean ± standard deviation. **p<0.01, ****p<0.0001. (G) Flow cytometry gating strategy for CD4 molecules.(H) Representative flow cytogram of CD4 molecules in different groups detected by flow cytometry. (I) Percentage of CD4 molecules in different groups. Data from each experimental group include at least three independent cell experiments, with results expressed as mean ± standard deviation. **p<0.01, ***p<0.001.

### 3. S protein interacts with mTOR, activates the mTOR-pS6 pathway, and promotes the activation of the latent reservoir

The mTOR pathway is closely associated with the pathogenic mechanisms of SARS-CoV-2. The S protein can activate the PI3K/Akt/mTOR pathway, regulating cellular responses, promoting viral replication, and inducing changes in cellular function. mTOR inhibitors effectively alleviate COVID-19 symptoms, further confirming their critical role.Additionally, mTOR plays a crucial role in HIV latent reservoir regulation and reverse transcription activation, influencing latent viral transcription through T cell activation signals and metabolic reprogramming. Therefore, mTOR not only regulates immune metabolism but may also participate in HIV latent reservoir activation. Given the central role of mTOR in both viruses, this study speculates that the S protein may indirectly alter the microenvironment of HIV host cells by activating the mTOR signaling pathway, thereby affecting the stability of the latent reservoir.

To preliminarily assess whether S protein regulates cellular function by activating the mTOR pathway, protein expression of mTOR and its phosphorylated forms (p-mTOR), pS6, and p-p70S6K, which are key molecules, was first detected by protein immunoblotting. The results showed that, in the presence of S protein, the expression levels of mTOR and p-mTOR significantly increased, and the expression of pS6 and p-p70S6K also increased, indicating that S protein can effectively activate the mTOR pathway (Figure 3A). These experimental results suggest that S protein may be closely related to the mTOR pathway in regulating cell function.

**Figure 3:**
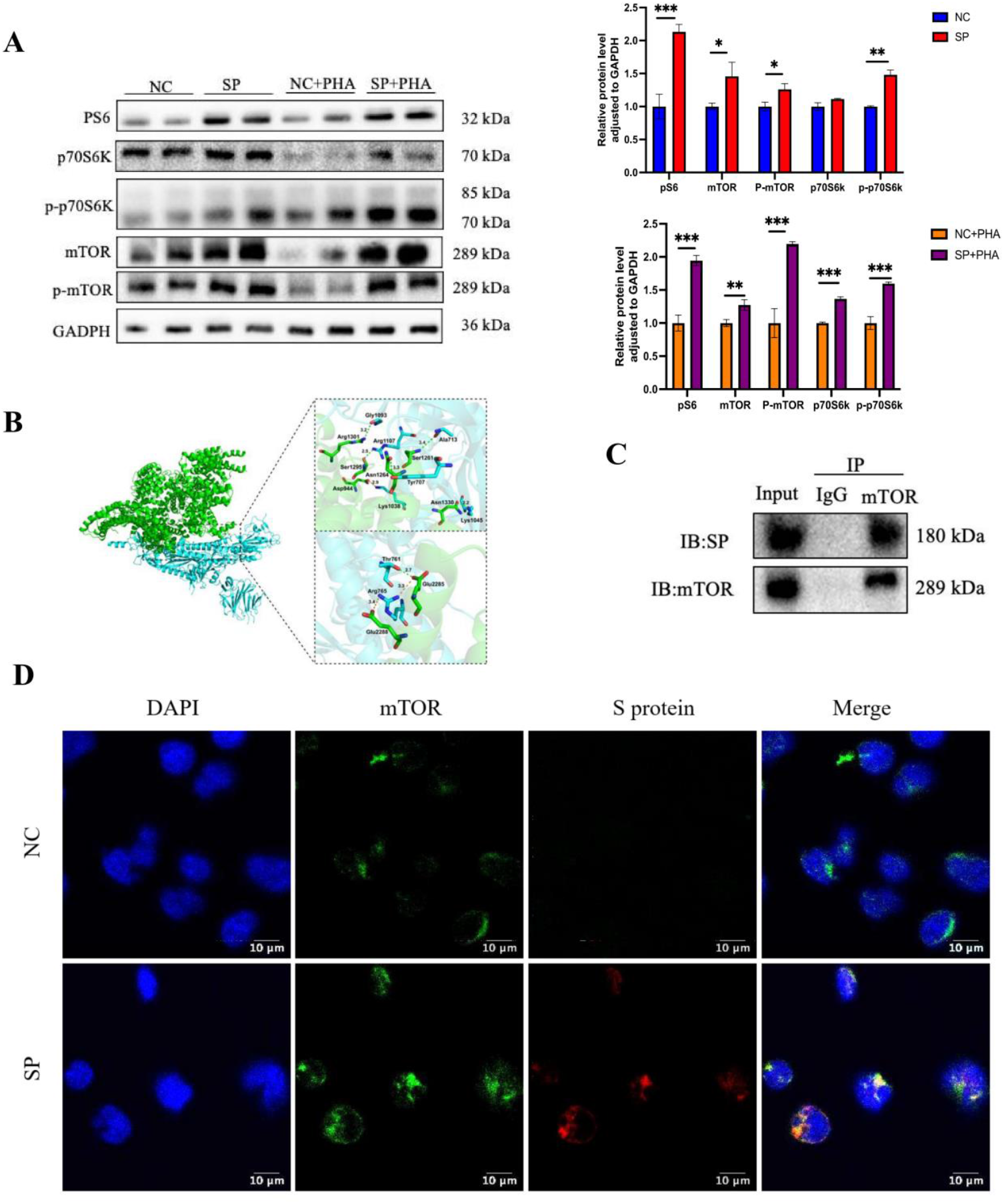
(A) Protein immunoblotting analysis of changes in mTOR, p-mTOR, pS6, p70S6K, and p-p70S6K protein levels in different groups. *p<0.05, **p<0.01, ***p<0.001. (B) Predicted interaction mode between mTOR and SPIKE protein via rigid docking.Green represents mTOR, blue represents SPIKE, green dashed lines indicate hydrogen bonds, and yellow dashed lines indicate salt bridges. (C) Immunoprecipitation validation of the interaction between S protein and mTOR, with Ig-g as the blank control. (D) Immunofluorescence colocalization validation of the interaction between S protein and mTOR.

Exploring the interaction between S protein and mTOR through molecular docking analysis. Upload mTOR and S protein to the zDOCK platform and predict their binding patterns using rigid docking. The results show that green dashed lines indicate hydrogen bonds, and yellow dashed lines indicate salt bridges. According to the z-score scoring standard, the optimal model scored −329.60, indicating that there may be a stable binding relationship between the two. mTOR forms hydrogen bond interactions with residues such as Arg 1301 and Ser 1295 of S protein through residues such as Gly 1093 and Arg 1107 (Figure 3B). This finding further confirms the possible binding between S protein and mTOR, providing a theoretical basis for subsequent experiments. To further validate the interaction between S protein and mTOR, CO-IP was performed in this study. It was found that S protein can form a complex with mTOR (Figure 3C). IFC also confirmed the conclusion that S protein interacts with mTOR. In Figure 3D, it can be clearly seen that the S protein of the SP group binds to mTOR, and the binding site after merging is shown as yellow fluorescence. These results confirm the binding relationship between S protein and mTOR, suggesting that S protein may play an important regulatory role in the cell through its interaction with mTOR, thereby regulating the activation process of the HIV latent reservoir. Based on the above experimental results, this study preliminarily infers that S protein may activate the mTOR pathway, thereby promoting the activation of the HIV latent reservoir. This finding provides strong experimental evidence for exploring the potential role of S protein in the activation of the HIV latent reservoir and offers important clues for further research into the mechanism of interaction between S protein and the mTOR pathway.

### 4. Inhibition of mTOR suppresses the activation of HIV by S protein

To validate whether S protein influences the activation of the HIV latent reservoir through the mTOR pathway, this study designed an experimental group treated with the mTOR inhibitor rapamycin (rapa). In this experiment, rapamycin was first added to the cell model for pretreatment, followed by the addition of PHA, to assess whether mTOR inhibitors could affect the activation of the mTOR pathway and the HIV latent reservoir triggered by S protein. The experimental groups established in this study included: the SP+PHA group (treated with S protein and PHA), the NC+PHA group (blank control group), the SP+PHA+rapa group (group treated with S protein, PHA, and rapa), and the NC+PHA+rapa group (blank cells treated with PHA and rapa).

To evaluate the effect of mTOR pathway inhibition on the HIV latent reservoir, the activation of the HIV latent reservoir in cells was detected by GFP fluorescence. As shown in Figure 4A, under a fluorescence inverted microscope, the fluorescence intensity of the NC+PHA group and the SP+PHA group was significantly reduced after the addition of rapa. Following this, GFP was quantified using flow cytometry, as shown in Figure 4BC, revealing that the number of GFP-positive cells in the NC+PHA group and SP+PHA group decreased significantly after the addition of rapa. Furthermore, we found that after the addition of rapa, the number of GFP-positive cells in the SP+PHA+rapa group remained higher than that in the NC+PHA+rapa group. RT-PCR analysis was performed to determine the transcription levels of LTR and GAG in each group. The results showed that in the SP+PHA+rapa group, the addition of rapamycin significantly suppressed the upregulation of Gag/LTR region transcription levels, indicating that mTOR inhibition has a significant effect on the activation of the HIV latent reservoir (Figure 4DE). Next, we detected the levels of P24 antigen using ELISA, and the results also showed that the P24 levels in the SP+PHA+rapa group were significantly lower than those in the SP+PHA group, further proving the role of the mTOR pathway in S protein-induced HIV latent reservoir activation (Figure 4F). Finally, flow cytometry analysis showed that CD4 molecule expression was reduced in the SP+PHA+rapa group compared with the SP+PHA group, further supporting the effect of mTOR inhibition on HIV latent reservoir activation (Figure 4H). In summary, the experimental results of rapamycin inhibiting the mTOR pathway indicate that mTOR plays a key role in the activation of the HIV latent reservoir induced by S protein, and that inhibiting the mTOR pathway can significantly suppress the activation of the latent reservoir.

**Figure 4:**
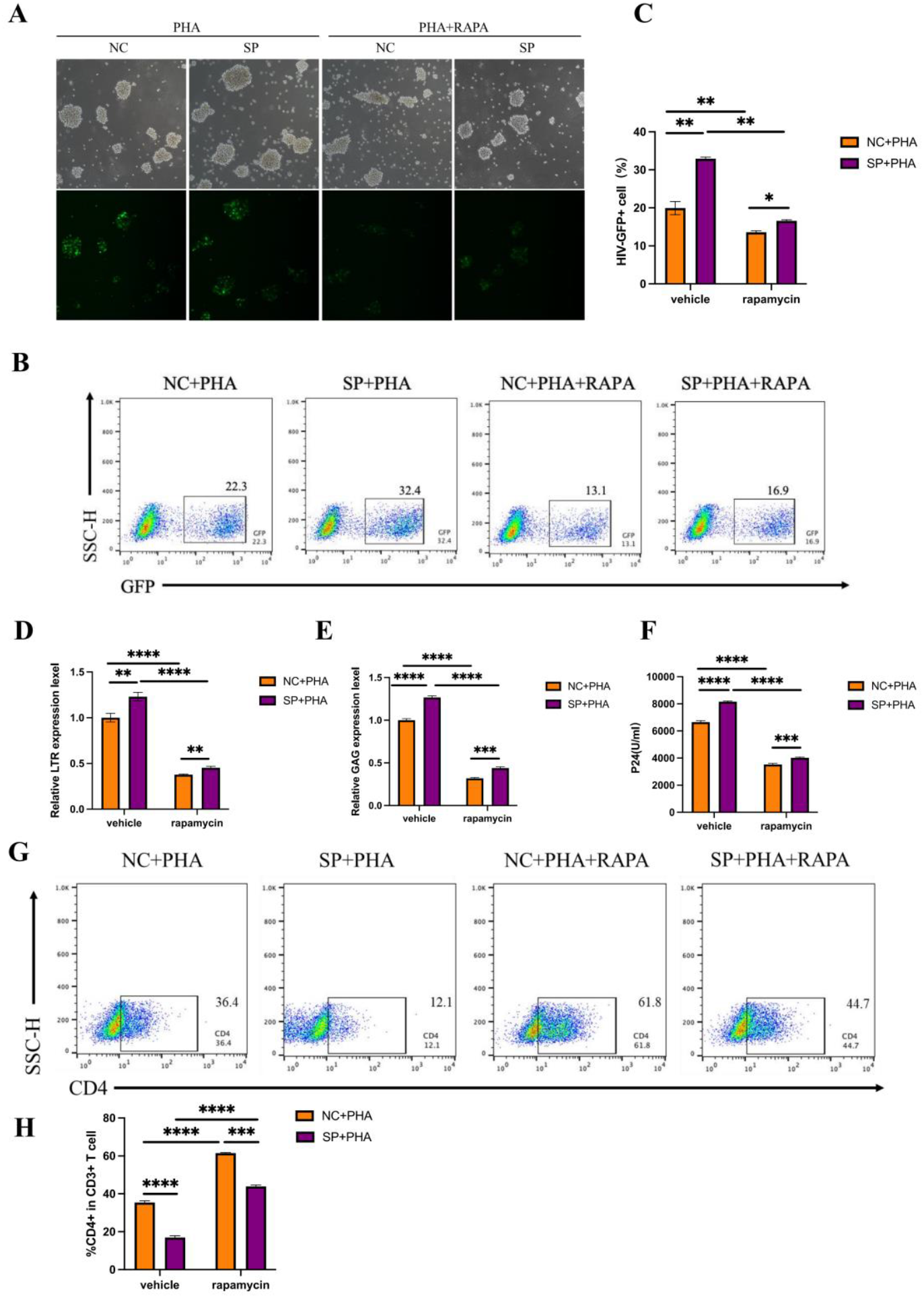
:(A) Electron microscope images of cells under different treatments showing GFP expression (magnification: 10×).(B) Representative flow cytometry plots of GFP-positive cells in different groups.(C) Statistical analysis of the percentage of GFP-positive cells. Data from each experimental group include at least three independent cell experiments, with results presented as mean ± standard deviation.**p<0.01. (D) Transcriptional levels of HIV LTR in different groups. Data from each experimental group include at least three independent cell experiments, with results expressed as mean ± standard deviation. **p<0.01, ****p<0.0001. (E) Transcriptional levels of HIV Gag in different groups. Data from each experimental group include at least three independent cell experiments, with results expressed as mean ± standard deviation. ****p<0.0001. (F) p24 antigen levels in the supernatant of different treatment groups. Data from each experimental group included at least three independent cell experiments, and results are expressed as mean ± standard deviation. ***p<0.001, ****p<0.0001. (G) Representative flow cytometric plots of CD4 molecules in different groups.(H) Percentage of CD4 molecules in different groups. Data from each experimental group included at least three independent cell experiments, and results are expressed as mean ± standard deviation. ***p<0.001, ****p<0.0001.

### 5. Replenishing mTOR reactivates the activation of s protein on HIV

On the basis of mTOR inhibition, we further investigated the effect of leucine repletion on the activation of the HIV latent reservoir by S protein through the mTOR pathway. Leucine is a known activator of mTOR. Following L-leucine replenishment, the proportion of GFP-positive cells recovered, suggesting that L-leucine can restore the activation of the HIV latent reservoir by S protein (Figure 5C). RT-PCR results showed that the addition of S protein significantly increased the transcription level of the HIV LTR/gag region, indicating that S protein promotes the activation of the HIV latent reservoir. Following the addition of the mTOR inhibitor rapamycin, HIV LTR transcription levels decreased significantly, indicating that inhibition of the mTOR pathway can attenuate the latent reservoir activation induced by the S protein. After adding leucine, the transcription levels of HIV LTR and Gag returned to levels close to those stimulated by S protein, suggesting that leucine supplementation may reactivate the HIV latent reservoir by restoring the activation of the mTOR pathway (Figure 5DE). Subsequently, we detected P24 secretion using the ELISA method, which also showed the same results (Figure 5F). Next, we used flow cytometry to detect changes in CD4 molecule expression in the replenishment experiment. The results showed that after the addition of leucine, the expression level of CD4 molecules in the cell model was replenished (Figure 5GH). In summary, these results further validate the important role of the mTOR pathway in S protein activation of the HIV latent reservoir and suggest that leucine may enhance the reactivation capacity of the latent reservoir by promoting the activation of the mTOR pathway.

**Figure 5:**
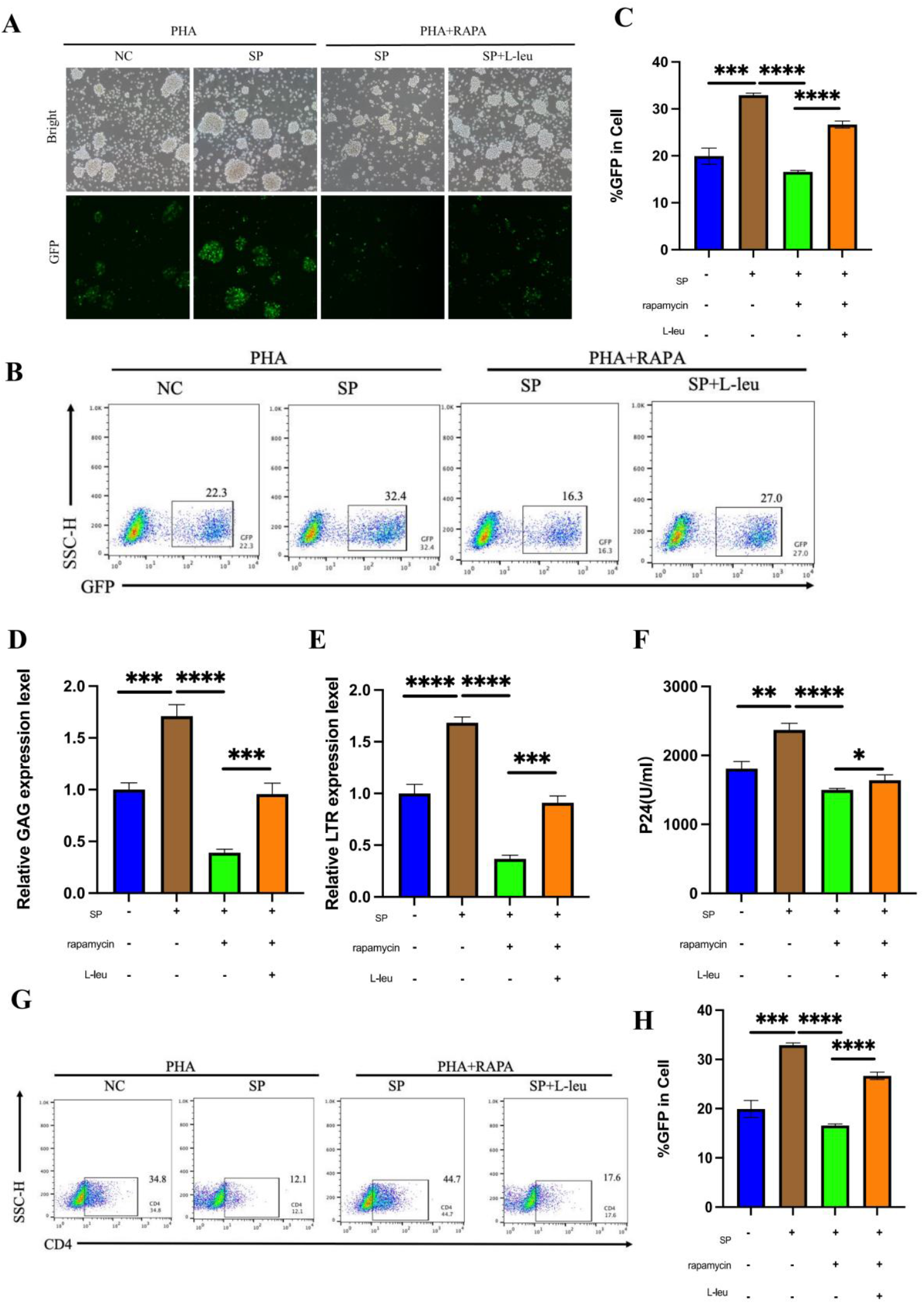
:(A) Electron microscope images of cells under different treatments showing GFP expression (magnification: 10×).(B) Representative flow cytometry plots of GFP-positive cells in different groups.(C) Statistical analysis of the percentage of GFP-positive cells. Data from each experimental group include at least three independent cell experiments, with results presented as mean ± standard deviation.***p<0.001, ****p<0.0001. (D) Transcriptional levels of HIV LTR in different groups. Data from each experimental group include at least three independent cell experiments, with results expressed as mean ± standard deviation.***p<0.001, ****p<0.0001. (E) Transcriptional levels of HIV Gag in different groups. Data from each experimental group include at least three independent cell experiments, with results expressed as mean ± standard deviation. ***p<0.001, ****p<0.0001. (F) Concentrations of p24 antigen in the supernatant of different treatment groups.Data from each experimental group include at least three independent cell experiments, with results expressed as mean ± standard deviation. *p<0.05, **p<0.01, ****p<0.0001. (G) Representative flow cytometric plots of CD4 molecules in different groups. (H) Percentage of CD4 molecules in different groups.Data from each experimental group include at least three independent cell experiments, with results expressed as mean ± standard deviation. ***p<0.001, ****p<0.0001.

### 6. S protein can initially activate the latent reservoir in ART patients

To investigate the potential role of the SARS-CoV-2 S protein in the HIV latent reservoir of patients receiving antiretroviral therapy (ART), peripheral blood mononuclear cells (PBMCs) were isolated from the peripheral blood of HIV-infected individuals who had been on long-term ART with sustained viral load suppression. Adeno-associated viral vector (AAV) was used to transiently transfect S protein into PBMC. CD3/CD28 magnetic beads were used to stimulate the cells, simulating the in vivo T cell activation microenvironment. To verify the expression of S protein in these cells, its expression was detected by RT-qPCR. The PCR results are shown in Figure 6A, confirming the successful transfection of S protein. First, we used PCR to detect viral expression in patients before and after treatment with S protein. Consistent with cell experiments, we found that S protein had a certain activating and amplifying effect on the latent reservoir of ART patients (Figure 6BC). To verify whether HIV viral particles activated by S protein possess infectivity, this study used the TZM-BL reporter cell line for in vitro infection experiments.

**Figure 6:**
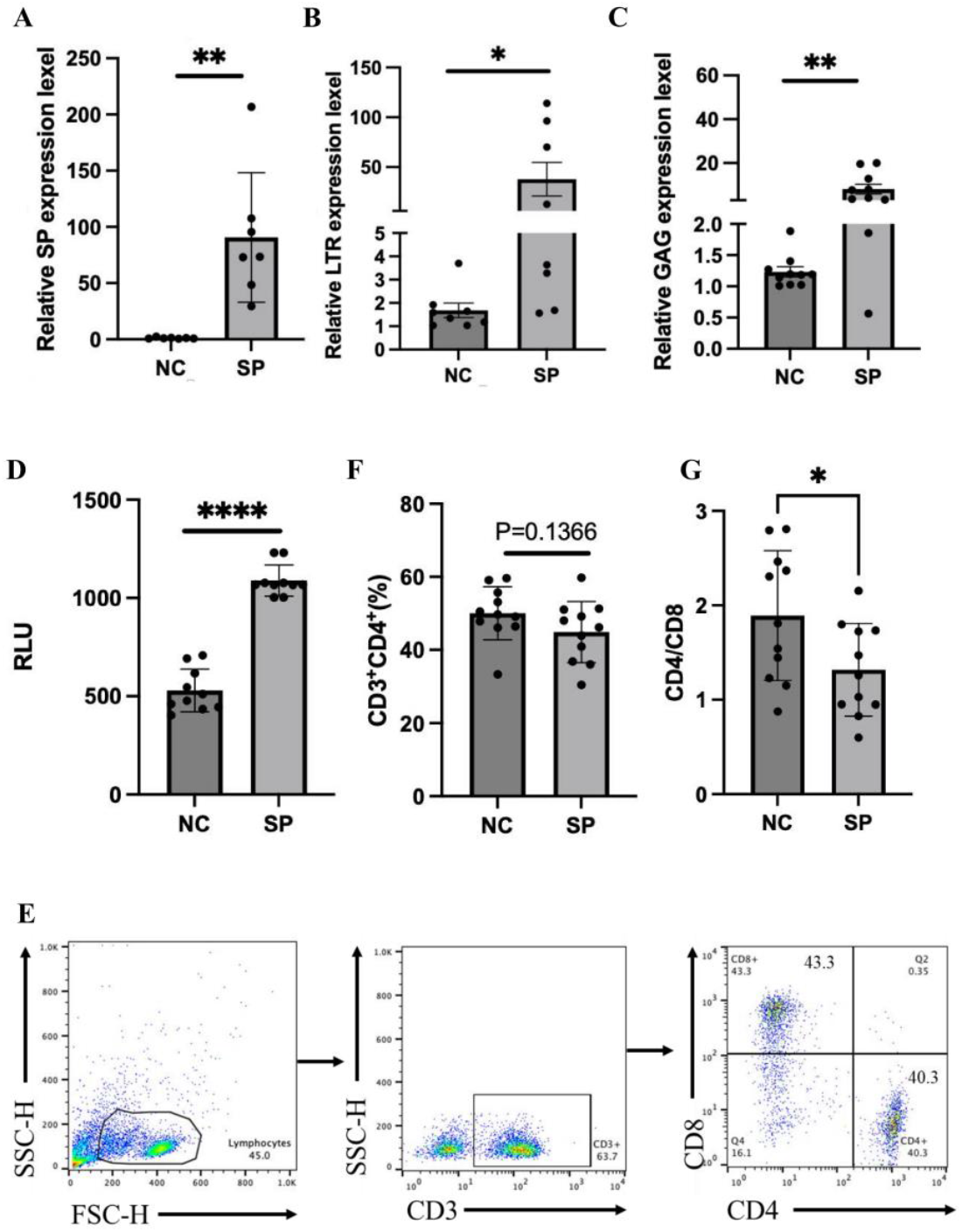
(A) Transcriptional levels of HIV S protein in PBMCs. Each point represents a patient. Results are expressed as mean ± standard deviation. **p<0.01 (B) Transcriptional levels of HIV LTR in PBMCs before and after S protein stimulation. Each point represents a patient. Results are expressed as mean ± standard deviation.*p<0.05 (C) Transcriptional levels of HIV Gag in PBMCs before and after S protein stimulation. Each point represents a patient. Results are expressed as mean ± standard deviation. **p<0.01 (D) HIV latent reservoir status in PBMCs before and after S protein stimulation, as measured by TZM-BL cells. RLU (relative luciferase expression) Each point represents a patient.Results are expressed as mean ± standard deviation. ****p<0.0001 (E) Representative flow cytogram of CD4 and CD8 cells in PBMC of patients detected by flow cytometry. (F) CD4 expression in patients before and after stimulation. Each point represents one patient. Results are expressed as mean ± standard deviation.*p<0.05, **p<0.01, ***p<0.001 (G) CD4/CD8 expression in patients before and after stimulation. Each point represents a patient. Results are expressed as mean ± standard deviation. *p<0.05, **p<0.01, ***p<0.001

The TZM-BL cell line is a widely used reporter gene cell line in HIV infection research. It integrates HIV LTR-driven luciferase and β-galactosidase genes, enabling indirect reflection of HIV viral infectivity through changes in fluorescence and enzyme activity. In the experiment, the supernatant of PBMCs treated with S protein was used to infect TZM-BL cells, and luciferase activity was detected after a certain period of time, with the infection level expressed as relative light units (RLU). The results are shown in Figure 6D. Compared with the control group, S protein promoted the activation of the HIV latent reservoir and enhanced the infectivity of virus release. Subsequently, we used flow cytometry to detect the proportion of CD4 cells in lymphocytes and the CD4/CD8 ratio in humans. We found that S protein can reduce the proportion of CD4 cells to a certain extent, which also indicates the activation of the HIV latent reservoir, although this finding was not statistically significant (Figure 6F). However, our comparison of CD4/CD8 cells before and after S protein stimulation revealed that S protein significantly reduced the CD4/CD8 ratio (Figure 5G). In summary, a series of experimental results indicate that S protein also promotes the activation of the HIV latent reservoir in ART patients, amplifying the activation effect of the latent reservoir. These findings provide strong evidence for the role of S protein in the activation of the HIV latent reservoir.

## Dicussion

The persistence of the HIV latent reservoir is a major obstacle to achieving a cure for AIDS, and the mechanisms underlying its reactivation are closely associated with the host immune microenvironment [26, 27]. In recent years, the disruption of the immune system caused by SARS-CoV-2 infection and its interaction with chronic viral diseases has attracted considerable attention. Clinical observations have shown that some HIV-infected individuals experience a transient rebound in viral load after contracting COVID-19, suggesting that SARS-CoV-2 may activate latent HIV through specific mechanisms [25]. However, the specific molecular mechanism underlying this phenomenon remains unclear.

This study aimed to investigate the effects of the S protein on HIV latent reservoir activation and its underlying pathway mechanisms. A cell line stably expressing the SARS-CoV-2 S protein was established using the classic HIV latent reservoir model J-Lat10.6 cells, and T cell activators PHA were used.This study revealed that the SARS-CoV-2 S protein promotes the activation of the HIV latent reservoir by activating the mTOR signaling pathway.Mechanistic studies have shown that the S protein can bind to mTOR, promote its phosphorylation, and activate the downstream pS6 signaling pathway, thereby driving the transcriptional activity of HIV long terminal repeats and ultimately promoting the activation of the HIV latent reservoir.This discovery not only reveals a new mechanism by which the S protein regulates the HIV latent reservoir, but also provides important clues for understanding the potential causes of viral load fluctuations in HIV-infected individuals following SARS-CoV-2 infection or vaccination.In vitro experiments showed that S protein treatment significantly increased HIV expression levels in latent infected cells, and this effect was specifically inhibited by the mTOR inhibitor rapamycin.Further validation of the key role of the mTOR pathway was achieved through replenishment experiments: L-leucine treatment restored the inhibitory effect of rapa on the HIV latent reservoir, indicating that mTOR activity is indispensable in S protein-mediated latent reservoir activation.

This study used the J-Lat 10.6 cell line as a model for the subsequent HIV latent reservoir. Its core advantage lies in the integration of a GFP reporting system based on HIV long terminal repeats [28, 29].Using methods such as flow cytometry, RT-PCR, and ELISA to detect the activation of HIV latent reservoirs.This study further characterized the latent reservoir model, which is similar to previous studies in that it does not express CD8 molecules, consistent with the characteristics of the HIV latent reservoir [30].

The mTOR signaling pathway plays an important role in cell metabolism, proliferation, and immune regulation.It was found that S protein activates the downstream pS6 signaling pathway by binding to mTOR and promoting its phosphorylation, thereby driving the transcriptional activity of HIV LTR.Introduction of PHA as a T cell activation stimulant. PHA, as a known latency reversal agent, can activate the TCR signaling pathway, induce transcription factors such as NF-κB and activator protein-1 (AP-1) to enter the nucleus, thereby breaking the latency state of HIV. This study found that after PHA stimulation, the GFP positivity rate, HIV transcript levels, and p24 release in the S protein-treated group showed significant synergistic enhancement effects, indicating that S protein not only independently activates the latent reservoir but also synergistically amplifies HIV transcription with PHA.

The activation of the mTOR pathway by the S protein is consistent with previous studies, and there have been multiple reports that SARS-CoV-2 can activate the mTOR pathway. In a proteomic study of the Huh7 liver cancer cell line, the EGFR family receptor tyrosine kinase (ErbB), hypoxia-inducible factor-1 (HIF-1), mTOR, and TNF signaling pathways were significantly upregulated during SARS-CoV-2 infection; These pathways share multiple key proteins, such as serine kinase (Akt), mTOR, eukaryotic translation initiation factor 4E-binding protein 1 (4E-BP1), and S6K1, whose phosphorylation levels significantly increase within 24 hours post-infection, indicating that SARS-CoV-2 activates the Akt-mTOR signaling pathway during the initial stage of infection [31]. Zhou et al. identified drugs such as the mTOR inhibitor rapamycin as potential treatments for COVID-19 through bioinformatics analysis of the HCoV-host interaction network [32]. Additionally, a recent study found that SARS-CoV-2 regulates host N6-methyladenine (m6A) modification by activating the mTORC1 signaling pathway to promote viral replication [33]. The novelty of this study lies in the fact that it is the first to experimentally validate the direct interaction between mTOR and the SARS-CoV-2 S protein, whereas previous studies have primarily focused on the indirect effects of free S protein on the cellular mTOR pathway. This finding provides more direct evidence for the mechanism by which the S protein regulates host signaling pathways. In addition, current mainstream COVID-19 vaccines primarily deliver the S protein via mRNA. This study utilizes a cell line stably expressing the S protein, enabling more accurate simulation of the sustained expression of the S protein following vaccine integration in vivo and its long-term impact on the HIV latent reservoir.This experimental design provides a model that more closely approximates physiological conditions for studying the sustained effects of S protein within cells and its regulatory mechanisms on the mTOR pathway.

The study found that inhibition of the mTOR pathway significantly reduced HIV expression, consistent with previous findings. The entry and replication of HIV-1 depend on the PI3K/Akt/mTOR signaling pathway. Specifically, HIV-1 envelope protein glycoprotein (gp120) binds to the CD4 receptor and activates the PI3K/Akt/mTOR pathway in the TZM-BL cell line; when phosphatase and tension protein homolog (PTEN), an inhibitor of the PI3K/Akt pathway, is overexpressed in target cells, HIV-1 entry is significantly blocked [34].

Additionally, in primary cell models, when HIV-1 reactivates from latency, the mTORC1 pathway is significantly upregulated in CD4^+^T cells [35]. The transcription and latency of HIV are also associated with mTOR regulation of metabolites required for epigenetic modifications, including acetyl-CoA and substrates of one-carbon metabolism. mTOR further regulates these metabolic processes by activating Akt and ATP citrate lyase [36], thereby influencing HIV latency and activation.

Currently, there are relatively few studies on the regulatory mechanism of S protein on HIV latent reservoirs in cellular systems. However, a recent study has shown that SARS-CoV-2 can reverse HIV latency in myeloid U1 cells either directly or through factors released by infected macrophages [37]. Unlike the subjects of this study, which focused primarily on myeloid cells, the present study focused on the latent reservoir in T cells [37]. Although the triggering mechanisms and research subjects differ, this finding is consistent with the experimental results of this study—that S protein promotes the activation of the HIV latent reservoir. This further supports the potential role of SARS-CoV-2 or its components in the regulation of the HIV latent reservoir.

In peripheral blood sample studies, HIV-infected individuals receiving ART had their PBMCs transfected with SARS-CoV-2 S protein in vitro, and significantly increased HIV proviral transcription activity and viral protein expression were observed. This finding reveals the activating effect of S protein on the HIV latent reservoir and suggests its potential regulatory role in the immune microenvironment of HIV-infected individuals. Consistent with the present study, previous research has observed that the BNT162b2 SARS-CoV-2 mRNA vaccine can induce HIV reactivation in ex vivo PBMCs from patients coinfected with SARS-CoV-2 and HIV, and this phenomenon is associated with the innate immune response to mRNA and the activation of NF-κB. In vivo experiments confirmed this hypothesis, showing that T cell responses associated with negative regulatory proteins (Nef) were significantly enhanced and further validated by granzyme B (gzm-B) enzyme-linked immunosorbent spot (ELISPOT) assays [38]. Additionally, a prospective open-label study found that in some patients who received SARS-CoV-2 mRNA vaccines, HIV viral load increased from <10 copies/mL to ≤40 copies/mL, accompanied by a mild decrease in CD4+ T cell counts [38]. However, some studies have also shown that COVID-19 vaccines do not affect viral load in ART patients; Among all HIV-positive patients who received the mRNA-1273 or BNT162b2 vaccine, although HIV RNA levels were detectable during the 6-month study period, only one participant had an HIV RNA level >200 copies/mL 6 months after the first vaccine dose [37].In a cohort study conducted in Hubei Province, China, HIV-positive participants experienced a statistically significant decrease in CD4^+^ T cell counts following the second vaccine dose; This phenomenon was observed after administration of the BNT162b2 mRNA vaccine and other vaccines recommended for HIV-infected individuals, consistent with the findings in this study that S protein introduction led to a reduction in CD4^+^cell counts [39].However, surprisingly, the study found that HIV viral loads decreased significantly in the cohort of HIV-infected individuals after vaccination; To investigate this phenomenon, researchers assessed the level of vaccine-induced T cell activation by detecting the expression of CD38 and human leukocyte antigen-DR (HLA-DR). The results showed a significant increase in the proportion of CD38^+^HLA-DR^+^ CD4^+^ T cells. The authors speculate that this decline in viral load may be attributable to ART or the elimination of activated HIV-1-infected CD4^+^ T cells by autologous cytotoxic T cells (CTLs) [39]. Based on the above, a possible dynamic mechanism is proposed: vaccination may induce a mild increase in HIV viral load in the short term, but as the proportion of cytotoxic T cells increases, viral load subsequently decreases [38, 39].

The interaction between SARS-CoV-2 infection and HIV latency is complex and multifaceted, with the mTOR pathway involved in cross-regulatory interactions across multiple pathways [40]. Future research should explore the following directions to deepen mechanistic understanding and translational research: First, the potential role of autophagy regulation in the latent reservoir. mTORC1 is a key inhibitor of autophagy, and autophagy can influence HIV latent reservoir stability by clearing viral proteins or regulating host metabolic states [17, 41]. Second, the association with metabolic reprogramming: whether mTOR activation provides energy support for latent reservoir activation by enhancing glycolysis (Warburg effect) [42]. Third, the hub role of inflammatory factor networks: the S protein can induce the release of pro-inflammatory factors such as TNF-α and IL-6 [43], which can influence the expression and activation of proteins like mTOR, activating the NF-κB pathway and directly driving HIV transcription [44].

## Conclusion

This study, by constructing an HIV latent reservoir model expressing the S protein, has for the first time demonstrated that the SARS-CoV-2 S protein can significantly promote HIV proviral transcription and viral particle release by directly binding to and activating the mTOR pathway.Mechanistically, the S protein remodels the host cell metabolic state by upregulating p-mTOR, pS6, and p-p70S6K levels, thereby releasing HIV latency. Intervention experiments targeting mTOR and validation results from PBMCs of ART-treated patients further support the central role of the mTOR pathway in S protein-mediated latency activation.This finding not only provides a molecular-level explanation for the interaction between HIV and COVID-19 but also lays the experimental foundation for personalized treatment strategies for co-infected patients such as prioritizing vaccination and monitoring viral rebound and targeted mTOR-based combination intervention strategies.

## Methods

### Statistical analysis

All data are presented as SEM ± mean. Statistical significance was determined using GraphPad Prism 8 (San Diego, California, USA). Two-sample unpaired Student’s t-tests were used for comparisons between groups. Differences were considered statistically significant: *p < 0.05, **p < 0.01, ***p < 0.001.

### Antibodies and reagents

Spike protein lentivirus (AD-SP), Spike protein adenovirus (LV-SP) purchased from HeSheng Gene Technology Co., Ltd. (Beijing, China). mTOR agonist L-leucine (HY-N0486) purchased from MCE; mTOR inhibitor rapamycin (AY 22989) purchased from MCE. PHA (L8754-5MG) purchased from Sigma.The anti-Spike antibody (40592-R493) for western blotting was obtained from SinoBiological; mTOR, phosphorylated mTOR (Ser2448), phosphorylated S6 ribosomal protein (Ser235/236), phosphorylated p70 S6 kinase (Thr389), and GADPH antibodies were purchased from Cell Signaling Technology;

### Patients

Peripheral blood mononuclear cells (PBMCs) used in this study were collected from HIV-1-infected individuals with confirmed HIV antibody positivity.Inclusion criteria include: receiving antiretroviral therapy (ART) for over 1 year, with HIV-1 RNA levels below the detection limit for at least 6 consecutive months, and peripheral blood CD4^+^ T cell counts greater than 350 cells/mL. Each donor provided 2 mL of peripheral blood for subsequent PBMC isolation and experimental research.All participants provided written informed consent. This study was approved by the Ethics Committee of Beijing Youan Hospital, Capital Medical University, and conducted in accordance with the Declaration of Helsinki (approval number: LL-2023-006-K). All participants provided informed consent, and the data used in this study are anonymous.

### Cell culture and cell line establishment

The HIV repository cell lines J-Lat Full Length Cells 10.6 (J-Lat 10.6) and TZM-BL used in this study were obtained from the National Institute of Allergy and Infectious Diseases (NIAID) of the National Institutes of Health (NIH) AIDS Reagent Program. Peripheral blood mononuclear cells (PBMCs) are obtained by separating human blood. J-Lat 10.6 cells and human PBMCs were cultured in RPMI 1640 complete medium containing 10% fetal bovine serum, 1% penicillin, and 1% streptomycin at 37°C in a culture incubator with 5% CO2 saturation and humidity.

Culture J-Lat10.6 in complete 1640 medium containing 10% FBS and 1% penicillin-streptomycin until the logarithmic growth phase.Subsequently, pre-prepared S protein lentivirus (MOI = 30) was added to the cells for infection, with an infection time of 12–24 hours. After infection, the viral medium was replaced with fresh culture medium, and the cells were continued to be cultured. Blasticidin (concentration: 5 μg/mL) was then added to initiate the selection of stably transfected cells, with a selection period of 3–7 days. The selected cells were confirmed by Western blot (WB) analysis to detect the expression of the reporter gene.Finally, expand the stably transfected cells and freeze them as needed for subsequent experiments. Isolate peripheral blood mononuclear cells (PBMCs) from whole blood and culture them in RPMI 1640 complete medium containing 10% fetal bovine serum, 1% penicillin, and 1% streptomycin at 37°C and 5% CO_2_ in a humidified incubator.Subsequently, perform transient transfection using adenovirus at an appropriate concentration. Add the virus to the cell suspension at the experimental design-specified viral dose, with a MOI of 30. After infection, continue culturing the cells for 24 hours to ensure adequate viral expression. Confirm successful adenovirus transfection and fluorescent gene expression via PCR. Proceed with subsequent experiments as required.

### Activation of the latent reservoir

First, culture J-LAT10.6 cells to the logarithmic growth phase. Next, treat the cells with PHA (final concentration of 5 μg/μL) to activate the HIV latent reservoir. The PHA treatment duration is 48 hours. After treatment, observe changes in GFP fluorescence in the cells using flow cytometry or fluorescence microscopy to assess the activation of the HIV latent reservoir.At this point, the treated cells can be used for subsequent experiments, sPBMCs from ART patients were cultured in RPMI 1640 medium containing 10% FBS and 1% penicillin-streptomycin, with cell concentration adjusted to approximately 1 × 10^6^ cells/mL. Following successful transfection of the S protein, CD3/CD28 magnetic beads (magnetic particles coated with CD3/CD28 antibodies) were added to the PBMCs at a 1:1 ratio, ensuring thorough mixing.To promote cell activation, IL-2 (final concentration 50 U/mL) was added to enhance T cell proliferation and activation. The cells were cultured in a 37°C, 5% CO_2_ incubator for 48 hours to activate HIV in the latent reservoir. Following activation, changes in HIV transcription and fluorescent labeling in the cells could be detected via flow cytometry to assess the activation status of the HIV latent reservoir.

### RT-PCR

Total RNA was extracted using a commercial kit, then transcribed into cDNA using the PrimeScript™ First Chain cDNA Synthesis Kit (Takara, China) according to the manufacturer’s instructions. Real-time PCR was performed using the Applied Biosystems 7300 Real-Time Fluorescent Quantitative PCR System (Thermo, USA) and TB Green® Premix Ex Taq™ (Takara, China). mRNA expression was normalized to GADPH expression.LTR-F:GCCTCCTAGCATTTCGTCACAT; LTR-R:GCTGCTTATATGTAGCATCTGAGG; Gag-F:GTCCAGAATGCGAACCCAGA; Gag-R: GTTACGTGCTGGCTCATTGC;GAPDH-F:CTCTGCTCCTCCTGTTCGAC;GAPDH-R:AGTTAAA AGCAGCCCTGGTGA.Spike-F:CTCTGCTCCTCCTGTTCGAC;Spike-R:ACGAGAACGGGACAAT CACC

### Flow cytometry

For the J-LAT10.6 cell line, prepare the cells in appropriate culture medium and perform surface labeling. Since the cells naturally express the GFP reporter gene, HIV latent reservoir activation is assessed by directly detecting GFP fluorescence using a flow cytometer. Additionally, J-LAT10.6 cells are labeled with CD3-percp and CD4-PE antibodies, incubated for 30 minutes at 4°C in the dark. After incubation, cells are washed twice with PBS to remove unbound antibodies. Then, data were collected using a flow cytometer, and appropriate channels were selected to detect GFP, CD3, and CD4 fluorescence signals. Data analysis was performed in FlowJo software to assess cell surface markers and GFP signals, and further analyze HIV latent reservoir activation and immune cell subset distribution. For human PBMCs, follow the above method using CD3-BV510, CD4-PE, and CD8-BV421 antibodies to label the PBMCs. Analyze the data in FlowJo software to assess cell surface markers and further analyze the activation status of the HIV latent reservoir and the distribution of immune cell subsets.

### ELISA

First, collect J-Lat 10.6 cells in the logarithmic growth phase, centrifuge them, adjust the cell concentration to 2 × 10^2^ cells/mL, and inoculate 2 mL per well into a 6-well plate. Incubate at 37°C and 5% CO_2_ for 48 hours. After 48 hours, collect the cell supernatant,centrifuge, and use the HIV-1 p24 antigen quantitative ELISA kit to detect antigen levels. The ELISA procedure is as follows: dilute the wash buffer, add the biotin reagent and sample or calibrator, and incubate for 60 minutes. Wash the plate five times, add the enzyme-labeled reagent, incubate for 30 minutes, and wash the plate five times again.Add substrate solutions A and B, mix thoroughly, and incubate for 15 minutes. Stop the reaction and measure the absorbance at 450 nm. Calculate the HIV p24 antigen concentration using the standard curve.

### Molecular docking

First, obtain the three-dimensional structures of MTOR and SPIKE protein and convert them into a file format supported by zDOCK. Then, upload MTOR and SPIKE protein to the zDOCK platform separately and set the rigid docking mode for molecular docking prediction. zDOCK will generate multiple complex models based on the rigid docking strategy. According to the obtained complex models, analyze their binding sites, binding energies, and interaction patterns.

### Luciferase assay

Resuspend the patient’s PBMCs in warm R10 medium, adjust the cell concentration to 2 × 10^2^ cells/mL, and seed into a 24-well plate at 1 mL per well. Add the corresponding treatment, gently tap to mix, and incubate at 37°C and 5% CO_2_ for 48 hours. After 48 hours, collect the cell suspension, centrifuge, and transfer the supernatant to a new tube.Next, digest TZM-bL cells with D10 medium, centrifuge, adjust the density to 0.2 × 10^2^ cells/mL, and seed into a 12-well plate with at least 8 wells. Add 200 μL of supernatant from each group. After 24 hours, aspirate the supernatant, add 200 μL of 1×Passive Lysis Buffer, and shake at room temperature for 15 minutes.Take 50 μL of the lysis buffer supernatant and add it to a 96-well plate. Add luciferase substrate, gently mix, and detect luciferase activity.

### Immunofluorescence colocalization assay for protein-protein interactions

Dilute CELL-TAK solution (5% acetic acid solution) with 0.1 M sodium bicarbonate buffer at a ratio of 1:30, then evenly dispense onto the surface of a small dish or coverslip. Incubate at room temperature or 37°C for 20 minutes to promote adsorption, followed by two gentle rinses with sterile water.Centrifuge 300 g of suspended cells for 5 minutes, resuspend in serum-free medium to an appropriate concentration (e.g., 1 × 10^2^ cells/mL), and add to the CELL-TAK-coated surface (50–100 µL per slide). Incubate at 37°C for 30–60 minutes to allow cells to adhere.Add 4% polyformaldehyde to fix for 10–15 minutes, rinse three times with PBS; then incubate with 0.1–0.5% Triton X-100 at room temperature for 5–10 minutes to permeabilize the cells, and rinse three times with PBS.Add blocking solution and incubate at room temperature for 30–60 minutes to reduce non-specific binding. Rinse off excess blocking solution with PBS, add diluted primary antibody, incubate at 37°C for 1–2 hours or overnight at 4°C, rinse three times with PBS, add secondary antibody, incubate at 37°C for 30–60 minutes, and rinse three times with PBS.Finally, add a mounting medium containing DAPI and other anti-fluorescence quenching agents, cover with a coverslip, and gently press. Observe protein colocalization in different fluorescence channels using an inverted fluorescence microscope.

### WB and co-IP

After lysing J-LAT cells, total protein was extracted and protein concentration was quantified using the BCA method. Equal amounts of protein samples were loaded onto a 12% Prosieve 50 gel for SDS-PAGE electrophoresis. After electrophoresis, proteins were transferred from the gel to a polyvinylidene difluoride (PVDF) membrane.After membrane transfer, the membrane was blocked with PBS containing 0.5% casein at room temperature for 1 hour. The membrane was then incubated with the primary antibody (1st antibody) in the blocking buffer at 4°C overnight. After three washes with PBS containing 0.05% Tween, the membrane was incubated with horseradish peroxidase-conjugated secondary antibody at room temperature for 1 hour.After washing extensively again, the membrane is incubated with Luminata Western HRP substrate, and chemiluminescence is used to detect protein expression. Results are captured using the Syngene G-box imaging system, and protein expression is quantified using Genesys software. GAPDH is used as an internal control protein for quantitative analysis. In the co-IP experiment, L-LAT cells are collected and lysed with lysis buffer, followed by centrifugation to obtain the supernatant.Incubate the supernatant with MTOR antibody for 2 hours, then incubate with Protein A/G magnetic beads at 4°C overnight to capture the MTOR-S protein complex. Wash the magnetic beads three times with PBS to remove unbound protein. Add SDS-PAGE loading buffer to dissociate the complex and perform SDS-PAGE electrophoresis.After transferring the membrane, detect with an S protein antibody and perform chemiluminescence detection using a secondary antibody. Use an IgG antibody as a negative control to ensure experimental specificity.

## Ethics approval and consent to participate

This study was approved by the Ethical Committee of Beijing Youan Hospital (Approval No. LL-2023-006-K). All participating patients provided informed consent, and the data used in the study were anonymized.

